# TOTEM: A web TOol for Tissue-EnrichMent analysis on gene lists

**DOI:** 10.1101/2024.04.04.588116

**Authors:** Veredas Coleto-Alcudia, Fidel Lozano-Elena, Gonzalo Vera, Isabel Betegón, Aditi Gupta, Idan Efroni, Ana I. Caño-Delgado

## Abstract

Analysis of spatiotemporal patterns of gene expression is crucial to decode biological systems responses. High-throughput sequencing allows in-depth transcriptome analyses and experimental designs, providing valuable reference expression atlases. Specifically, testing overrepresentations of tissue-specific genes based on these atlases can provide valuable insights; however, such an approach is not accessible to inexperienced users. Here, we introduce TOTEM (<u>TO</u>ol for <u>T</u>issue-<u>E</u>nrich<u>M</u>ent), a web tool designed to calculate enrichment values per tissue by identifying tissue-specific genes from an organ/organism of interest given a user gene list. Results are visually represented, and user’s gene classified. The utility of TOTEM is manifest when using integrated single cell expression atlases, enabling the study of complicated tissues, with the maximum possible resolution. Its effectiveness is validated by the study of BRL3 role in stress specifically from the vascular tissues. Finally, TOTEM’s modular design allows for continual integration of new experiments. TOTEM can be freely accessed at: https://totemwebtool.com.

## 1. Introduction

The importance of compartmentalization in organisms is manifest (Banani et al., 2017; Bar-Peled & Kory, 2022). The discovery of tissue-specific and cell-specific functions has advanced the understanding of tailored response mechanisms and disease causes, as well as guiding drug design and delivery strategies to effectively tackle these issues (Hekselman & Yeger-Lotem, 2020; Kiyotani et al., 2021; Pfeiffer et al., 2022; Zhao et al., 2020). However, the study of tissue-specific effects of treatments and/or mutations is not straightforward unless they are disrupted and/or isolated from their natural context. In this aspect, massive gene expression data (i.e., *RNA-seq*) has traditionally been a powerful tool for investigating the effects and causes of impaired gene function and treatment effects. In particular, the recent adoption of single-cell high-throughput sequencing (*scRNAseq*) has allowed the transcriptional profiling of individual cells in a massive way, theoretically reaching the maximum resolution possible at the organism level (Elmentaite et al., 2022; McFaline-Figueroa et al., 2020). However, these approaches have particular drawbacks: in the bulk *RNAseq* approach, the elucidation of tissue-specific responses or transcriptional profiles relies on the physical separation of tissues (e.g., tissue disruption, FACS, laser microdissection) and comparing transcriptional responses between them (Byron et al., 2016; Han et al., 2015; Stark et al., 2019). In contrast, in the *scRNA-seq* approach, the identification of transcriptionally similar cell groups relies on clustering algorithms. Yet, to unequivocally assign tissue identities to these groups of cells, it is mostly relied on transcriptional atlases validated through other methods (Kiselev et al., 2019). Further, when profiling treated and/or diseased cells, these might change their “natural” transcription fingerprints, possibly leading to an incorrect cell clustering that masks the tissue-specific responses (Cuperus, 2022; Kharchenko, 2021; Li & Wang, 2021; Ren et al., 2018).

In this context, the use of canonical gene expression atlases can be a powerful resource. Unbiased gene expression profiling (i.e. validated, wild type and control conditions) has yielded detailed expression maps at organ or tissue levels that integrate spatial and/or temporal information of the organ or organism under study (Brady et al., 2007; Ganini et al., 2021; Haniffa et al., 2021). These large datasets constitute valuable references, which are often not fully exploited. We propose to leverage all the information gathered in these cell expression atlases to increase the biological insights provided by *omics* experiments by adding a new functional layer (rather than just topological), including the tissue- or cell-specific origin of transcriptional responses. This can be, for example, testing overrepresentations of tissue-specific features in a given gene set and/or intersecting these with lists of tissue-specific genes with the aim of identifying key genetic players.

However, the manipulation of such large datasets can be complicated, especially for non-experienced users. Indeed, tools that facilitate visualization and management of gene expression in a spatial context are available and widely used, such as the multipurpose tool Genevestigator (Hruz et al., 2008), Expression Atlas (Moreno et al., 2022), plant-specific eFP browser (Sullivan et al., 2019; Winter et al., 2007), and Arabidopsis root specific with single ell resolution RootCellAtlas (Mironova, 2022). Although these available tools have their own strengths regarding gene expression topology, these either lack high spatial resolution or only allow for single gene query or gene pair comparisons.

With TOTEM webtool, we aim to provide a simple tool for tissue enrichment analysis based on gene lists and tissue-specific expression derived from existing datasets. With a particular focus on plants, we integrate all available Arabidopsis *scRNAseq* maps in a canonical atlas whose tissues have been identified using experimentally validated tissue markers. This integrated atlas exemplifies what can be done on other species of agronomical interest. From this, multiple functional characterization analyses might be derived (e.g., *pseudo-temporal* reconstruction, Gene Ontology Enrichment Analysis (GEOA), Pathway Enrichment) based on tissue-specific subsets identified by TOTEM. Although TOTEM is specifically developed for plants, analogous approaches may be used for other organisms, such as mammals and insects, using the same web tool.

## 2. Design

With increasing evidence pointing towards the huge importance of plant tissue-specific responses to drive adaptation and specialized functions (Fàbregas et al., 2018; Kajala et al., 2021; S. Kim et al., 2020), we identified the lack of holistic tools to explore the tissue contributions in the observed transcriptional responses. In previous works, we used literature-available expression maps to intersect our genes of interest (i.e. Differentially Expressed Genes, DEGs) with previously identified tissue-specific genes and test eventual tissue enrichments (Fàbregas et al., 2018). The fruitful outcomes from this approach prompted us to implement it in a simple and user-friendly web tool that also allows the visualization of enrichment results and link them to further functional characterization steps.

Current tools designed for similar approaches, as TissueEnrich (Jain & Tuteja, 2019), TEnGExA (Rawal et al., 2021) and LaminaRGeneVis (E. H. Kim et al., 2022), require of bioinformatic skills and/or are limited to mammals. Whereas available tools focused to visualization purposes, such as Genevestigator (Hruz et al., 2008), Expression Atlas (Moreno et al., 2022), eFP browser (Sullivan et al., 2019; Winter et al., 2007) or RootCellAtlas (Mironova, 2022) are limited to single gene query or pair comparisons (Supplementary Table 1). In addition, single-cell RNA sequencing is bringing unprecedented insight into tissue-specific responses (Cuperus, 2022; Fahlgren et al., 2023; McFaline-Figueroa et al., 2020) and there exist a wealth of transcriptional maps in plants, mainly in the model plant Arabidopsis (Cuperus, 2022; Efroni et al., 2016; Minne et al., 2021; Shahan et al., 2021; Shaw et al., 2021) but also on crops like maize (Marand et al., 2021) or rice (Zhang et al., 2021). Visualization tools specific for these experiments are available too (Ma et al., 2020; Wendrich et al., 2020). However, in these particular experiments and visualizers, not all tissues are represented, and some have limited resolution (i.e., the number of cells captured). Therefore, we sought to integrate these individual datasets to cover the maximum number of tissues and with the maximum possible resolution, thereby creating a canonical expression atlas that would be the reference to be used as background for enrichment testing in the approaches implemented in TOTEM.

## 3. Results

### TOTEM: A web tool for tissue enrichment analysis in plants

TOTEM is a user-friendly web tool designed to provide spatiotemporal information on gene lists and contribute to their functional characterization. Its main function is to calculate enrichment values per tissue based on a user-provided gene list and a reference gene expression atlas. TOTEM also represents the enrichment values in an organ/organism colored diagram, facilitating the interpretation, pinpointing which genes are specific to a particular tissue, and allowing further functional characterization.

TOTEM input page has two drop-down tabs with the available species and experiments (Fig 1A, input 1(i1)) and a description box where experiment information is displayed upon selection along with an uncolored organ/organism diagram. Importantly, a blank text field is left (with default ID examples) for users to paste their list of gene identifiers of interest (Fig 1A, i2) and a button to run enrichment once all variables are selected. The results page is composed by barplot depicting the actual enrichment scores (Fig 1B, output 1(o1)) and a colored diagram representing such scores (Fig 1B, o2), both downloadable. In addition, a drop-down tab allows users to explore which genes in their list are specifically expressed in a particular tissue and send them for functional characterization (Fig 1C). The functional characterization page displays detailed information on selected tissue-specific genes (based on Phytozome annotation data files (Goodstein et al., 2012), Fig 1C, o3) and allows running Gene Ontology Enrichment Analysis (GOEA) and KEGG pathway enrichment analysis, illustrating outcomes in a graph (Fig 1C, o4). In addition, an additional tab is opened, which allows mapping of the expression of specific genes across all the atlas in case a single cell experiment was selected (Fig 1D, o5).

**Figure 1.**
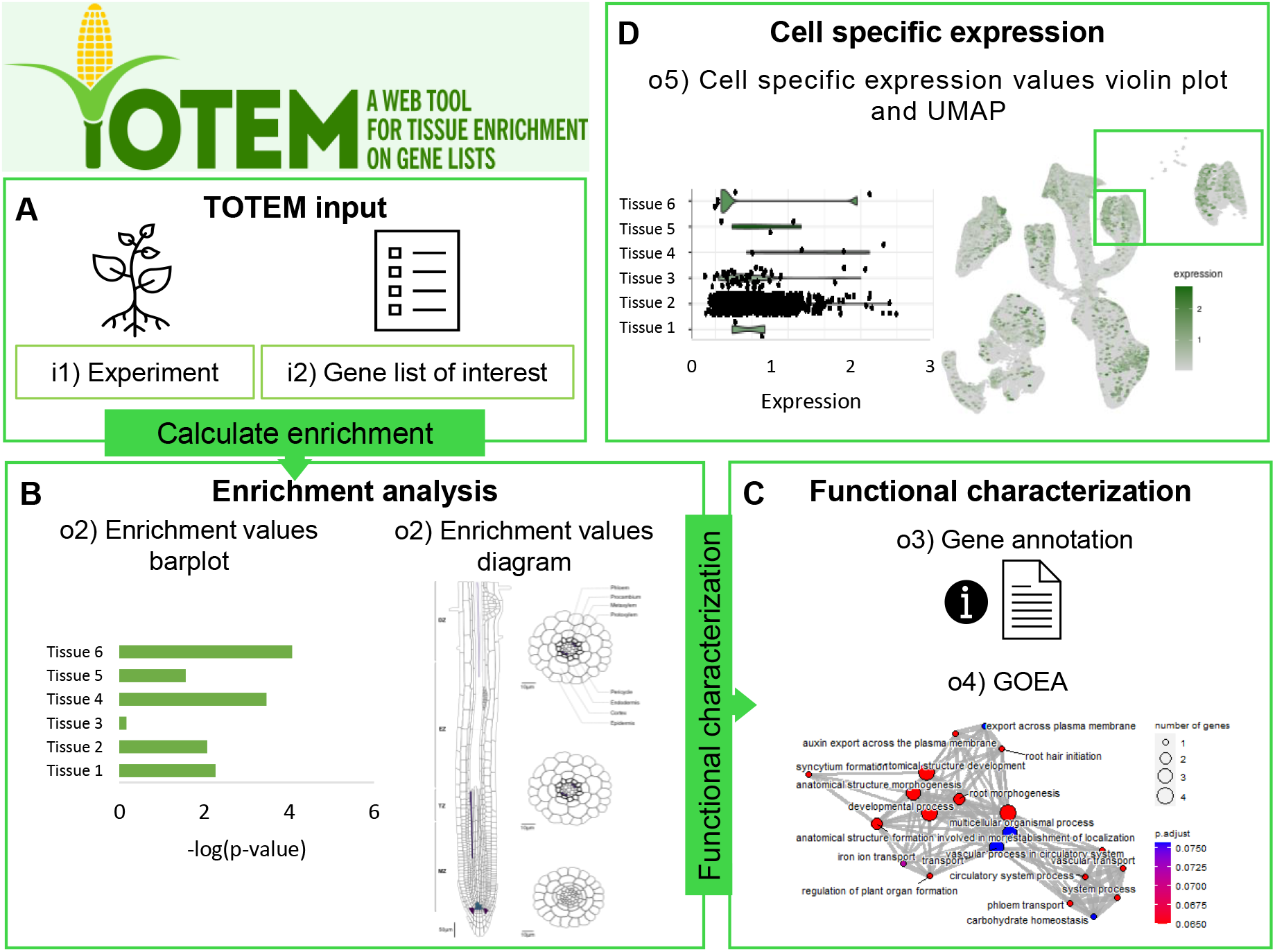
Summary of TOTEM workflow and its basic inputs, processes and outputs. **A)** TOTEM input page displays two drop-down tabs to select the specie and the experiment to be used as reference dataset (input1, i1) for further steps. When the experiment is selected, its corresponding description and blank SVG image are prompted. An example of gene identifiers corresponding to the selected experiment are displayed in a text field where user can paste their gene list of interest (input2, i2). **B)** Upon submission, the program calculates enrichment scores across tissues and represents them in the Enrichment Analysis result page in form of barplot (output 1, o1). The returned p-values are used to color the vectorial image of the experiment selected (output 2, o2). Finally, intersections are performed between user gene list (i2) and a list of tissue-enriched genes determined in the experiment (i1). Upon tissue selection, genes from the user list that are enriched in a particular tissue are shown in a text box. **C)** To better characterize the genes enriched in interesting tissues, a table with information about the genes (output 3, o3), as well as a GO enrichment and KEGG pathway enrichment analysis are available in Functional Characterization page (output 4, o4). **D)** If the selected experiment is a single cell one, the expression of a selected gene from the user list can be checked in each single cell of the atlas (output 5, o5).

It should be kept in mind that the generated image (Fig 1B, o2) is just a visual representation (may not depict all tissues examined in the experiment) and that the threshold used to consider the input list enriched in genes specific of a particular tissue is an arbitrary decision. TOTEM aim is to provide biological insights based on gene lists and published spatiotemporal datasets; therefore, users might be more astringent or loose with their threshold depending on their objectives. TOTEM is entirely written in R-Shiny language (Chang et al., 2023) and designed in a modular way, so new experiments can be added without major code rearrangements. The code is freely available at (https://github.com/CRAGENOMICA/totem) according to the GNU-LGPL v2.1 license.

### Bulk RNAseq & scRNAseq atlases

Classic transcriptomic experiments (e.g., microarray) can also be included in TOTEM, provided that lists with tissue-specific genes can be identified. Indeed, these are often underexploited valuable assets. For, example we included the first available spatio-temporal map of Arabidopsis primary root (Brady et al., 2007), which used a validated tissue isolation approach and physical sectioning of the organ. To illustrate the use of TOTEM, we analyze results derived from Differential Expression Analysis of a previous study. Upregulated genes due to the overexpression of one member of the Arabidopsis brassinosteroid (BRs) receptor family, BRL3, under drought conditions (Fàbregas et al., 2018) were compared to the radial patterns of the Arabidopsis root spatiotemporal map (Brady et al., 2007). Enrichment of genes specific to the phloem, pericycle, and lateral root primordia was found among the DEGs (Fig. 2A), which match with the native expression pattern of BRL3 (Fàbregas et al., 2013) and the vascular predominance on the overexpressor lines (Fàbregas et al., 2018).

**Figure 2.**
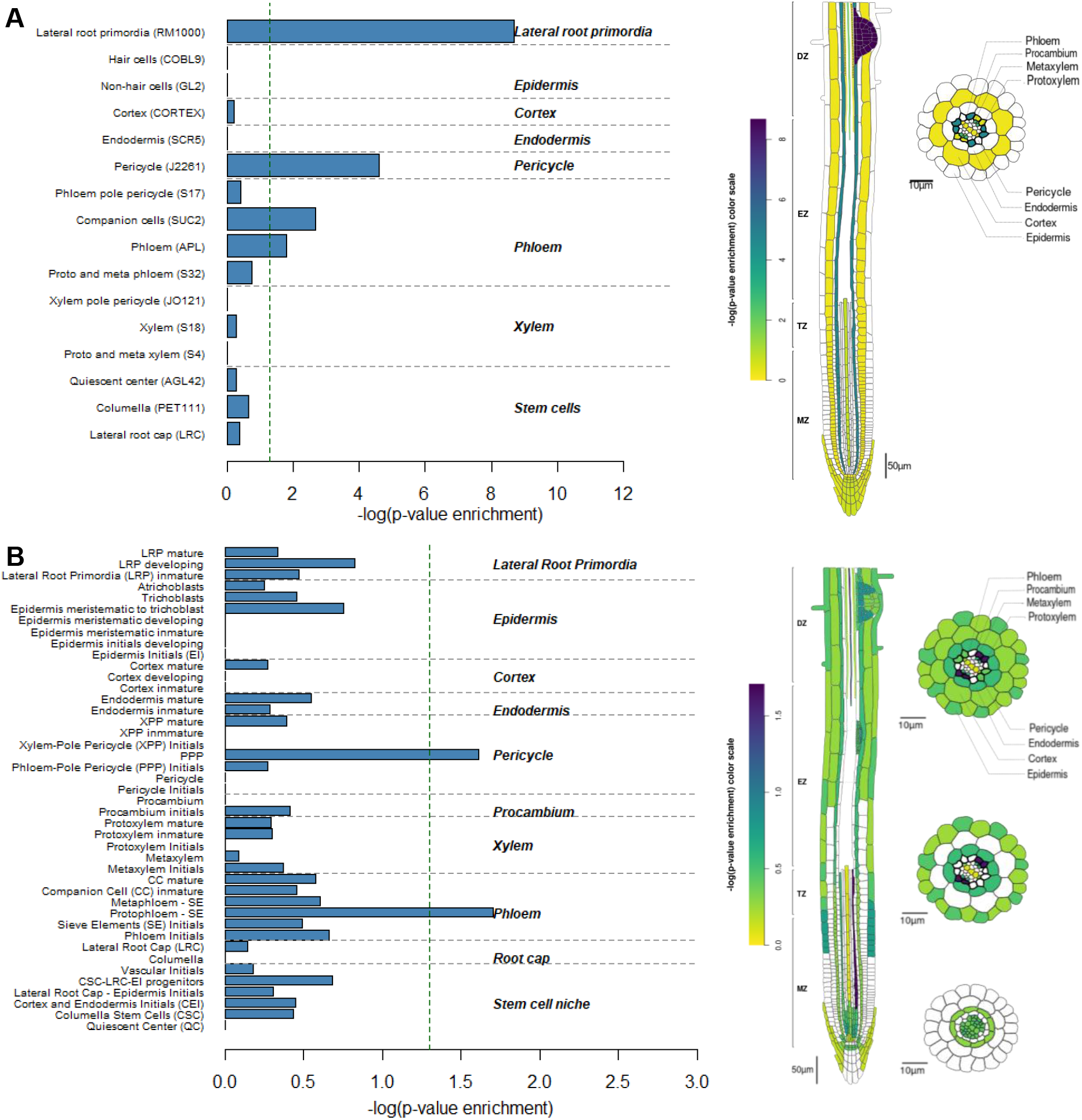
Validation of TOTEM with bulk transcriptomic data. Diagram outputs from the result page for Arabidopsis root, based on **A)** radial patterns of Brady et al., 2007 and **B)** single-cell atlas. Query genes are upregulated (FDR<0.05 and |logFC|>1.0) under drought conditions, derived from differential expression analysis of *BRL3ox* vs. Col-0 roots (Fàbregas et al., 2018). The dashed green vertical line of the barplot corresponds to p-value = 0.05.

If a single-cell experiment is selected, TOTEM performs the same approach but using the scRNAseq reference atlas as background. To illustrate this, we selected the integrated canonical atlas of Arabidopsis primary roots (see next section) and input the upregulated genes of BRL3ox plants (Fàbregas et al., 2018). The results revealed similar enrichment patterns compared to microarray experiments, but with an increased resolution, pointing towards more specific cell populations (Fig. 2B).

The functional characterization module can rapidly provide information on user genes specifically expressed in a particular tissue. For example, in our dataset, unveiling the metabolic relevance of vascular-specific DEGs in specialized sugar metabolism, which is in accordance with previous results (Fig. S1, (Fàbregas et al., 2018)). Furthermore, intersecting the tissue-specific gene lists of the two dimensions considered in the experiment (radial and longitudinal root sections, Brady et al., 2007) also allows the identification of very specific genes and if there is a very prominent enrichment in a particular group (Fig. S2).

To illustrate the use of other organisms and organs, drought responsive genes that correlated with developmental defects in a BR-deficient tomato cultivar (Lee et al., 2018) were tested against the tomato fruit development and ripening expression map (Shinozaki et al., 2018). The results showed an over-representation of genes specific to the columella and placenta during the early phases of fruit maturation (Fig. S3), which is consistent with the role of BR in development.

### Implementation of an integrated single cell atlas of Arabidopsis

To offer the highest resolution possible in Arabidopsis root and leaf organs, TOTEM integrates the different available scRNAseq datasets in a canonical reference expression atlas. For Arabidopsis roots, we integrated a total of 154005 cells (Wild type, control conditions) coming from six available studies (Denyer et al., 2019; Jean-Baptiste et al., 2019; Ryu et al., 2019; Shahan et al., 2022; Shulse et al., 2019; Wendrich et al., 2020). The integration resulted in root single-cell expression atlas of >150k high quality cells, spanning 26672 genes (See methods, Fig 3A-B, Fig S5A, Supplementary Table 2). To identify cell types, we used a mixture of standard clustering algorithms, the Garnett classifier (Pliner et al., 2019), and experimentally validated genetic markers (Fig. S5B-C, Supplementary Table 3; see Methods). In order to gain resolution and include a “temporal” dimension, we segmented the cell types into smaller clusters according to developmental trajectories in a pseudo-time analysis (Fig. S5D-F, See methods). A total of 43 cell populations were identified (Fig. 3B, Fig S5G). These included specific plant root tissues at different developmental stages, as protoxylem initials, immature and mature cells, procambium or immature, developing, and mature cortex. The identification of key gene markers based on these tissues or cell populations (See methods) constitute the background for the enrichment tests run by TOTEM main feature. In the case of Arabidopsis leaf, the construction of such canonical expression atlas follows an identical approach (Fig. 3A, Fig. S6) (J.-Y. Kim et al., 2021; Lopez-Anido et al., 2021), however this is limited by the number of studies available and cells sequenced and influenced by the intentional biases introduced on these studies (Supplementary Table 2, Supplementary Table 4). From the integration of Arabidopsis leaves, scRNA-seq resulted in an expression atlas of 11413 high quality cells spanning 22773 genes and clustered in 34 cell populations (Fig. 3C, alt. Fig S6G).

**Figure 3.**
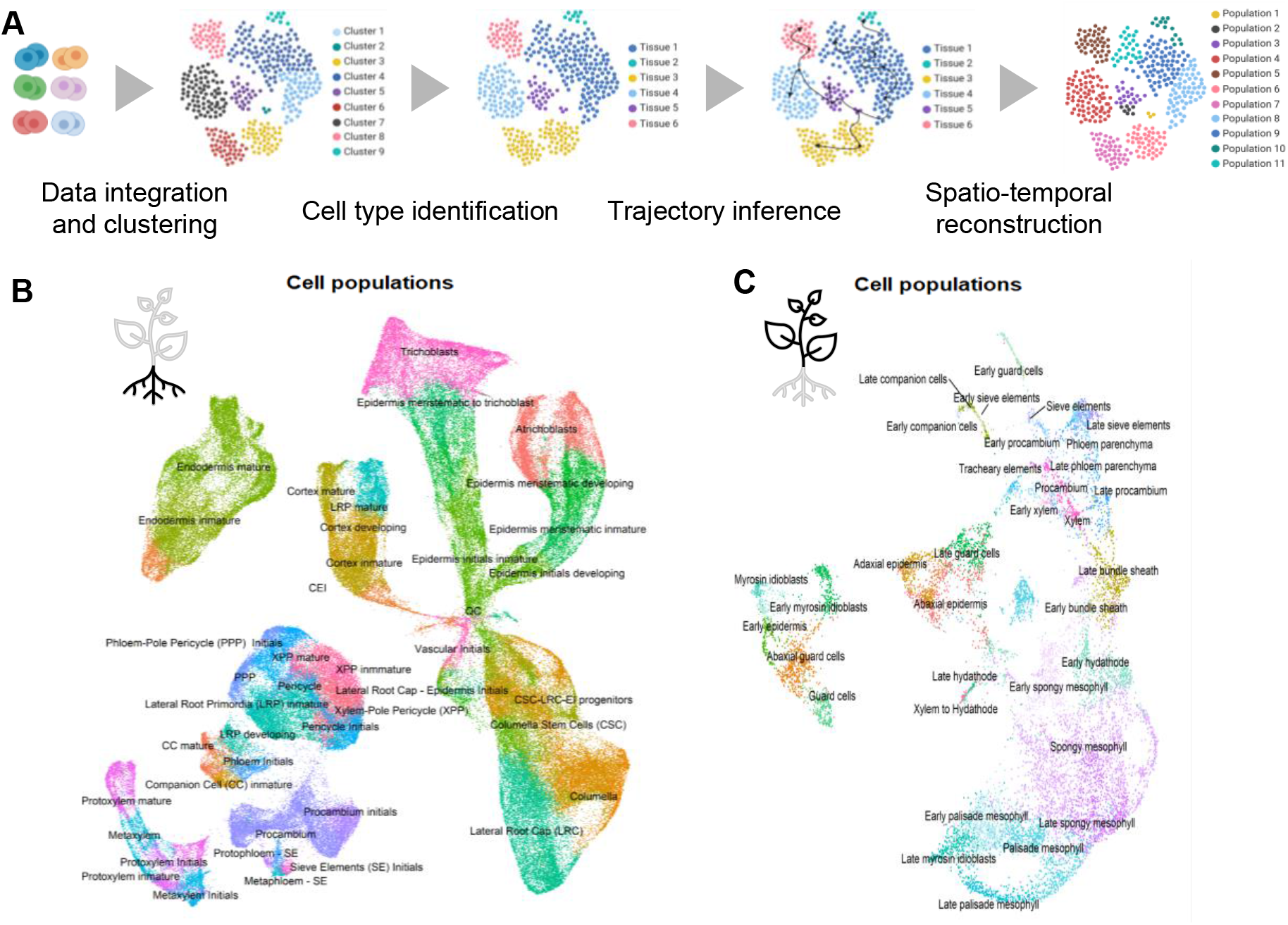
Integration of scRNAseq experiments in a canonical atlas. **A)** Overview of scRNA-seq dataset integration into a unique atlas. Cell populations identified in the Arabidopsis **B)** root and **C)** leaf single-cell expression atlas.

### Molecular signatures of BRL3 in phloem with single cell resolution

To demonstrate a potential application of the tool, we used TOTEM to analyze the results emanating from a single-cell sequencing coupled with Fluorescence-Activated Cell Sorting (FACS). In this experiment, the role of BRL3 receptor in phloem cells was investigated. We drove BRL3 expression specifically in the phloem using the companion-cell-specific promoter SUC2 (Imlau et al., 1999) and tagged the receptor with GFP (pSUC2:BRL3-GFP plants, Fig 4A), provoking tissue-specific overexpression (Gupta et al., 2023). The transgene was introduced into Arabidopsis wild-type plants (*Col-0* ecotype) and we used 5-day-old transgenic roots to isolate GFP positive cells through FACS (see methods). The mRNA of 153 GFP-positive cells was sequenced, yielding 102 high-quality cells after filtering, which were subjected to further analysis. We annotated these cells on the integrated single-cell Arabidopsis root atlas and, unsurprisingly, a high proportion of cells were annotated over companion cells and vascular stem cells (Fig 4B, Fig S4A), matching the expression pattern of SUC2 (Fig 4A). This result showcases the potential of TOTEM to be used as validation tool: It maps the user’s experiment against an integrated version of several spatiotemporal expression atlas. We then sought to analyze the transcriptomic changes induced by increased BRL3 expression in the phloem. So, we run Differential Expression Analysis between our GFP-positive cells (BRL3 overexpression) and cells from the annotated cell populations of the reference integrated single cell atlas (Fig 4B), in analogy to previous approaches (Butler et al., 2018; Li et al., 2022; Lotfollahi et al., 2023; Salcher et al., 2022). Based on the 2121 DEGs, we used TOTEM to map the transcription response (putatively, most of them should affect phloem) and to identify tissue-specific genes. Independently of the Arabidopsis experiment used as background, the obtained enrichment patterns are similar, showing enrichment in phloem, lateral root primordia and pericycle genes among upregulated DEGs (Fig 4C, Fig S4B) whereas enrichment among downregulated genes were in stem cells and epidermis (Fig. 4E). These findings suggest that BRL3 in phloem cells promotes vascular functions related to responses to stress (transport, defense, and stress responses GO terms, Fig. 4D, S4C-D, Supplementary Table 5), while repressing functions associated with stem cell niche and outer root layer development (regulation of translation and gene expression GO terms, Fig. 4F, Supplementary Table 6). Altogether, these outcomes exemplify the new topological information layer and refreshed the point of view that TOTEM can provide when interpreting *omics* analyses.

**Figure 4.**
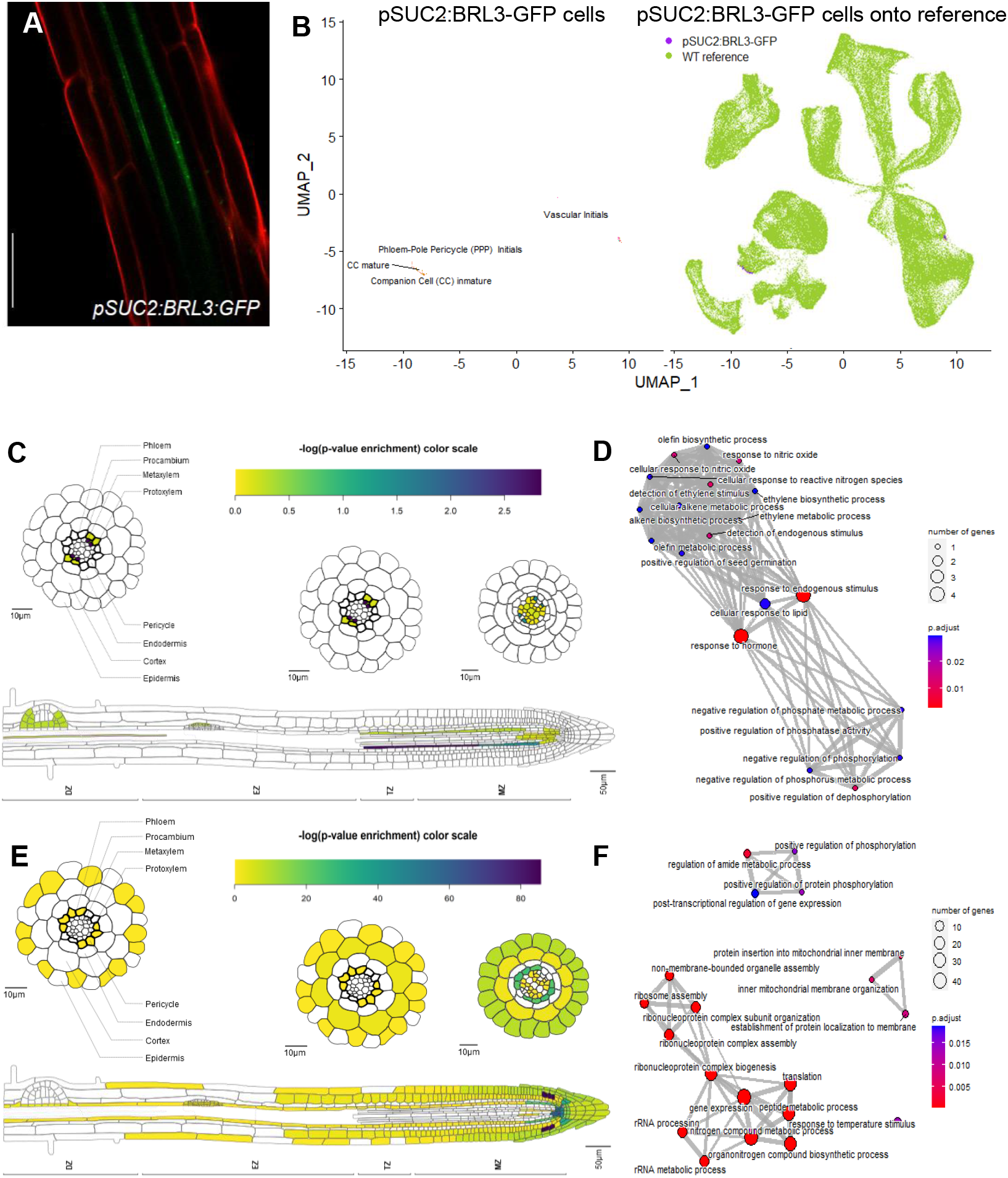
pSUC2:BRL3-GFP cells pattern enrichment with TOTEM. **A)** BRL3 expression in Arabidopsis root phloem cells under SUC2 promoter. **B)** pSUC2:BRL3-GFP cells (purple) annotated over single-cell root TOTEM reference dataset (green). TOTEM enrichment pattern in single-cell experiment of deregulated genes (FDR<0.05, |log2(FC)| >0.5) in the pairwise comparison of pSUC2:BRL3-GFP cells vs. WT cells of TOTEM reference atlas of **C)** upregulated genes and **E)** downregulated genes. GO term enrichment of upregulated genes in **D)** companion cells immature and downregulated genes in **F)** vascular initials cell populations are depicted. The size of the dots corresponds to the number of DEGs enriched in the GO term. The color of the dots corresponds to the adjusted p-value (FDR). The dashed green vertical line of the barplot correspond to p-value = 0.05.

## 4. Discussion

The development of TOTEM was motivated by the willingness to further exploit published spatio-temporal expression datasets and the necessity of additional functional annotation approaches to characterize gene lists, for example, those derived from differential expression analysis. To our knowledge, there is no available tool capable of dealing with gene lists that yield statistical values and visual representations (Supplementary Table 1). Now, we implement the tissue-enrichment approach in an easy and user-friendly web tool. Although simple, TOTEM can rapidly add a topological information layer to the group of genes of interest. This might be very meaningful for users aiming to understand gene regulatory networks in a spatial context or decipher tissue-specific responses. Because the TOTEM experiment library is easily expandable, new experiments and species can be easily incorporated upon request.

### 4.1. Limitations

The main function of TOTEM is a simple enrichment approach coupled to graphical representation, which has no major limitations *per se*. Thus, TOTEM limitations are associated with the dependence on published/available experiments. We identified three limitations that could influence the accuracy of the outputs. These are, from less to more influential:

i. The number of experiments used as background. For example, if TOTEM outputs are based on a single experiment (rather than on a combination or an integrated experiment, e.g., the single-cell atlas of Arabidopsis roots), the exposure to potential biases from the original study increases. To mitigate this, we recommend that the user carefully read the description provided and, if necessary, refer to the original study from which the atlas serves as background.
ii. The correct identification of cell populations. This should be taken into account, especially in experiments where the identification of cell populations is “unbiased” and/or based on clustering algorithms (such as single-cell experiments). These are very sensitive to inputs parameters and offer a mathematical solution that does not necessarily reflect the biology underlying. This is not a problem for well-known and differentiated tissues; however, when a user is seeking very specific cell populations, this may become a limitation. This is not a limitation in experiments where tissue isolation or identification have been empirical (e.g. FACS, microdissection).
iii. The identification of marker genes of each tissue (which will be the basis for the enrichment test) is very sensitive to astringency parameters and identification algorithms used, and is therefore sensitive to false positives. Additionally, the size of these gene markers lists use to be large (depending on the astringency of the analysis) leading to large enrichment values. These although “numerically” significant, it may not be biologically significant. Therefore, we advise users to have a holistic view and compare across all tissues tested.

TOTEM tool is intended to provide with additional biological insights to the classical functional characterization steps. We warn potential users that TOTEM does not provide categorical results. It provides insights that may be subjected to interpretation. So, we recommend validating results before initiating further works uniquely based on TOTEM outputs.

## Acknowledgments

The authors acknowledge all members of the Caño-Delgado laboratory for critical testing of the web tool and their feedback on its implementation. A.I.C.-D. has received funding from the European Research Council (ERC) under the European Union’s Horizon 2020 research and innovation programme (grant agreement No 683163). A.I.C-D. is a recipient of a grant (FEDER-BIO2016-78150-P) funded by the Spanish Ministry of Economy and Competitiveness-National Research Agency, and the European Regional Development Fund. V.C-A is funded by the Severo Ochoa PhD Fellowship PRE2019-088780 funded by MCIN/AEI/ 10.13039/501100011033 and by “ESF Investing in your future”. F.L.E. and A.G. have received funding from ERC-2015-CoG–683163 granted to the A.I.C.-D. laboratory. A.G. received funding from the Severo Ochoa postdoctoral fellowship. We acknowledge financial support from the Spanish Ministry of Economy and Competitiveness, through the “Severo Ochoa Programme for Centres of Excellence in R&D” 2016-2019 (SEV-2015-0533)” and by the CERCA Programme / Generalitat de Catalunya.

## Author contributions

A.C.D. and F.L.E. conceived the project. F.L.E., V.C.A. and G.V. implemented the web tool and introduced the data. A.G. and I.B. generated the transgenic plants, designed, and performed tissue-isolation and *scRNAseq* experiments. I.B., I.E. and V.C.A. analyzed the *scRNAseq* data. F.L.E., V.C.A., and A.C. wrote the manuscript.

## STAR Methods

### TOTEM INTERFACE

The TOTEM interface is written in Shiny R and is partially based on previous approaches from collaborators (Rafael-Palou et al., 2012). Its core function translates numeric values into color intensities for graph objects in a Scalable Vector Graph (SVG) file. The remaining operations are performed in core R, which is connected to the TOTEM interface. The architecture of the web tool separates each dataset into a separate R-data file that is loaded when the experiment is selected by the user. Each Rdata includes: (i) an R list in which each entry is a vector (tissue) of genes specifically expressed in a single tissue; (ii) the gene universe of the experiment (detected genes in RNAseq and single-cell RNAseq or probes in microarrays) that will serve as background for the enrichment function; (iii) a custom function for each experiment that represents the enrichment values in the form of a barplot (a dashed green line is depicted at p-value 0.05); and (iv) a custom function for each experiment that translates enrichment values to scaled scores to for the SVG drawing and solves eventual incompatibilities with the SVG file (i.e., domains that overlap in some regions).

### DATA ACQUISITION AND ANALYSIS

#### Data acquisition

To construct the root and leaf atlas, we selected available scRNA-seq datasets that contained cells representing WT main tissues of the organs, six datasets in the case of Arabidopsis roots, and two in the case of Arabidopsis leaves (Supplementary Table 2). For bulk experiments, microarray and RNAseq datasets of Arabidopsis, Sorghum and tomato were retrieved from (Brady et al., 2007; Davidson et al., 2012; Shinozaki et al., 2018), respectively.

#### Construction of single cell RNAseq atlases

Integration of root and leaf single-cell datasets was performed in R (version 4.1.2) using Seurat (version 4.1.0, (Hao et al., 2021)), filtering out cells with less than 200 expressed genes and cells with more than 10% of their reads mapped to mitochondrial and chloroplast genes. Also, genes expressed in less than 3 cells were filtered out. The 2000 features that had high cell-to-cell variation in the datasets were extracted from all root and leaf datasets, except for the Shahan dataset, in which 6000 were extracted owing to their large size. Root datasets have a higher conservation of cell types because of the homogeneity of samples, but this is not the case for leaf datasets. Taking this into account, we followed the recommendations of the Seurat team in their vignette “Fast integration using reciprocal PCA (RPCA)” (https://satijalab.org/seurat/articles/integration_rpca.html) and log-normalized root datasets and integrated them using canonical correlation analysis (CCA) and normalized leaf datasets with SCT transformation and integrated them using reciprocal PCA (RPCA). After integration and scaling of the atlas in the case of the root, a PCA analysis was performed using 50 PCs, the same PCs as in the UMAP reduction for visualization. A resolution value of 0.8 was used in clustering of both atlases. The integration resulted in an scRNAseq root atlas of 154005 high quality cells, 26672 genes and 37 clusters (Fig 3A, S5A-B) and an scRNAseq leaf atlas of 11413 high-quality cells, 22773 genes and 22 clusters (Fig 3B, S6A-B).

Cell type identification of the clusters was performed by checking the expression of experimentally tested marker genes of the roots and leaves in the literature (Supplementary Tables 3 and 4). The same list of known marker genes was used then to automatically confirm the cluster identity using Garnett tool (Pliner et al., 2019). 20 cell types were identified in the root atlas (Fig S5C) and 11 in the leaf atlas (Fig. S6C). Next, we studied developmental progression using trajectory inference within the cell types. We followed the same two approaches for root and leaf datasets. First, we used CytoTRACE (R package version 0.3.3, (Gulati et al., 2020)) in the complete atlas to assign a pseudotime value related to the differentiation state of cells (Fig S5D, S6D). Then, we divided the atlas into cell groups based on their cell type identity using Dyno (R package version 0.1.2, (Saelens et al., 2019)) to infer trajectory groups using the ten cells with the highest CytoTRACE pseudotime values as initial cells and the slingshot method as the optimal method based on this prior knowledge in all cell groups and both datasets (Fig. S5E, S6E). Trajectory analysis revealed us 95 and 51 trajectory groups in root and leaf atlases, respectively (Fig S5F, S6F). Finally, we grouped the trajectories according to their cell identities. This resulted in a spatiotemporal reconstruction of the atlases to identify all the cell populations of roots and leaves in terms of both cell type and developmental stage identities. From the 95 trajectory groups, we were able to identify 43 cell populations in different developmental stages in root atlas (Fig 3B, S5G) and from the 51 trajectory groups, 34 cell populations were identified in leaf atlas (Fig 3C, S6G). The high number of cell populations identified in the integrated atlas can address the limitations of using individual scRNA-seq datasets, as different cell types and stages of development are present in the atlas.

As in bulk experiments of TOTEM, we must define the tissue-specific genes. For this, we took advantage of the Seurat function “FindMarkers” to find markers of each cell population via differential expression analysis. We defined as cell population markers those genes that are differentially expressed (p-value adjusted by Bonferroni < 0.01; average log fold change |FC| > 1) in one cell population compared to all other cell populations using Wilcox method. Specifically, a marker gene is the one which is expressed in more than 25% cells in the complete atlas, but also the percentage of cells expressing the marker in the cell population is the double than in the rest of the cell populations.

#### pSUC2 single cell RNAseq library preparation and sequencing

For pSUC2:BRL3-GFP single-cell experiment, we utilized the mcSCRB protocol, adapted from (Bagnoli et al., 2018). For protoplasting, roots from pSUC2:BRL3-GFP expressing plants (Gupta et al., 2023) were cut and shaken in protoplasting solution for 20 minutes to separate the meristems. These were then transferred to another well containing 50µl of protoplasting solution and shaken for 1 hour. The plant material was then filtered using a 40 µm strainer. The protoplasting solution was prepared by adding 1.5% Cellulase, 0.5% Macerozyme, 0.4M Mannitol, 20.48 mM MES and 0.02M KCl; adjusting pH to 5.7; heating to 55° C and cooling to room temperature; and adding 0.2% BSA and 0.02M CaCl2.

For Fluorescence-Activated Cell Sorting (FACS), protoplasts underwent sorting using a high-speed cell sorter. Wild type protoplasts were utilized to establish a gate for GFP-negative cells, after which GFP positive samples were sorted to obtain positive cells. Single cells were sorted into 96-well DNA LoBind plates (Eppendorf) containing 5 µl of lysis buffer, comprising a 1:500 dilution of Phusion HF buffer (New England Biolabs), 1.25 µg/µl Proteinase K (Clontech), and 0.4 M barcoded oligo-dT primer (E3V6NEXT, IDT). Following sorting, plates were promptly spun down and frozen at −80°C.

For cDNA synthesis, the 96-well plates were incubated at 50°C for 10 minutes to digest proteins, followed by heat-inactivation of Proteinase K at 80°C for 10 minutes. A 5 µl reverse transcription master mix was dispensed per well, containing 20 units Maxima H-enzyme (Thermo Fisher), 2x Maxima H-Buffer (Thermo Fisher), 2 mM each dNTPs (Thermo Fisher), 4 µM template-switching oligo (IDT), and 15% PEG 8000 (Sigma-Aldrich). cDNA synthesis and template-switching were conducted for 90 minutes at 42°C. Barcoded cDNA was pooled and cleaned-up using SPRI beads, then purified cDNA was eluted and residual primers digested. Following heat-inactivation, PCR amplification was performed using a PCR master mix, and samples were purified again using SPRI beads. Quality control was assessed by PCR, and cDNA was quantified using the Quant-iT PicoGreen dsDNA Assay Kit (Thermo Fisher).

Library preparation was conducted using the Nextera XT DNA Library Prep kit (Illumina) with modifications, starting from 1 ng of preamplified cDNA and adjusting reagent volumes. Libraries were checked for size and quality using a DNA High Sensitivity Bioanalyzer chip (Agilent) and quantified with the Quant-iT PicoGreen dsDNA Assay Kit (Thermo Fisher). Pooled libraries were equimolarized, selecting the main peak, and further confirmed via agarose gel electrophoresis. After band extraction, library quality was reassessed, and libraries were paired-end sequenced on high output flow cells of an Illumina HiSeq 1500 instrument.

#### pSUC2 single cell analysis

After sequencing pSUC2:BRL3-GFP cells, the FASTQ files underwent a filtering process for poly-A reads utilizing a customized script and were then aligned to the TAIR10 genome using STAR 2.7.1. The read calling process was performed using the zUMI pipeline developed by Parekh et al. (2018). Transcripts were extended at their 3’ by 1Kb (or until closest gene) to capture reads originating from misannotated 3’UTRs. Bioinformatic analysis were performed in R (version 4.1.3) using Seurat package (Hao et al., 2021) to construct Seurat object by keeping cells with unique features (genes) higher than 400 and less than 7500, cells that have number of total molecules (counts) lower than 100000, cells which percentage of mitochondrial and chloroplast genes lower than 10 % and genes detected in at least 3 cells. pSUC2:BRL3-GFP cell raw counts from Seurat object were annotated over the integrated Arabidopsis thaliana single cell atlas root with SingleR package (Aran et al., 2019) using the final cell populations as labels. Predicted labels of pSUC2:BRL3-GFP cells (pSUC2 dataset hereafter) were transferred to its Seurat object to merge with the integrated Arabidopsis thaliana single-cell atlas root. As described in (Hao et al., 2021), a new UMAP reduction was computed after merging the pSUC2 dataset and the integrated atlas. Differential expression analysis was carried out using the Seurat function “FindMarkers” between pSUC2 cells and the integrated atlas cells that belong to the cell populations where pSUC2 cells were annotated. Genes differentially expressed (p.value adjusted by Bonferroni < 0.05; average log fold change |FC| > 1) were used as input for enrichment analysis using TOTEM web tool.

### DATA AND CODE AVAILABILITY

TOTEM can be freely accessed in https://totemwebtool.com. The source code is deposited in https://github.com/CRAGENOMICA/totem and can be freely reused and modified according to the GNU-LGPL v2.1 license.

## Supplemental information titles and legends

**Supplementary Table 1.** Feature comparison of TOTEM with other available gene expression visualizers (alphabetical order).

**Supplementary Table 2.** Overview of the root and leaf scRNA-seq datasets used for integration and atlas construction.

**Supplementary Table 3.** Experimentally validated gene markers used for cell type identification in the Arabidopsis root scRNA-seq atlas.

**Supplementary Table 4.** Experimentally validated gene markers used for cell type identification in the Arabidopsis leaf scRNA-seq atlas.

**Supplementary Table 5.** Description of the upregulated genes of pSUC2:BRL3-GFP vs. WT reference cells enriched in cell populations in the TOTEM single-cell experiment.

**Supplementary Table 6.** Description of the downregulated genes of pSUC2:BRL3-GFP vs. WT reference cells enriched in cell populations in the TOTEM single-cell experiment.

