## Supplementary figures for "TOTEM: A web TOol for Tissue-EnrichMent analysis on gene lists"

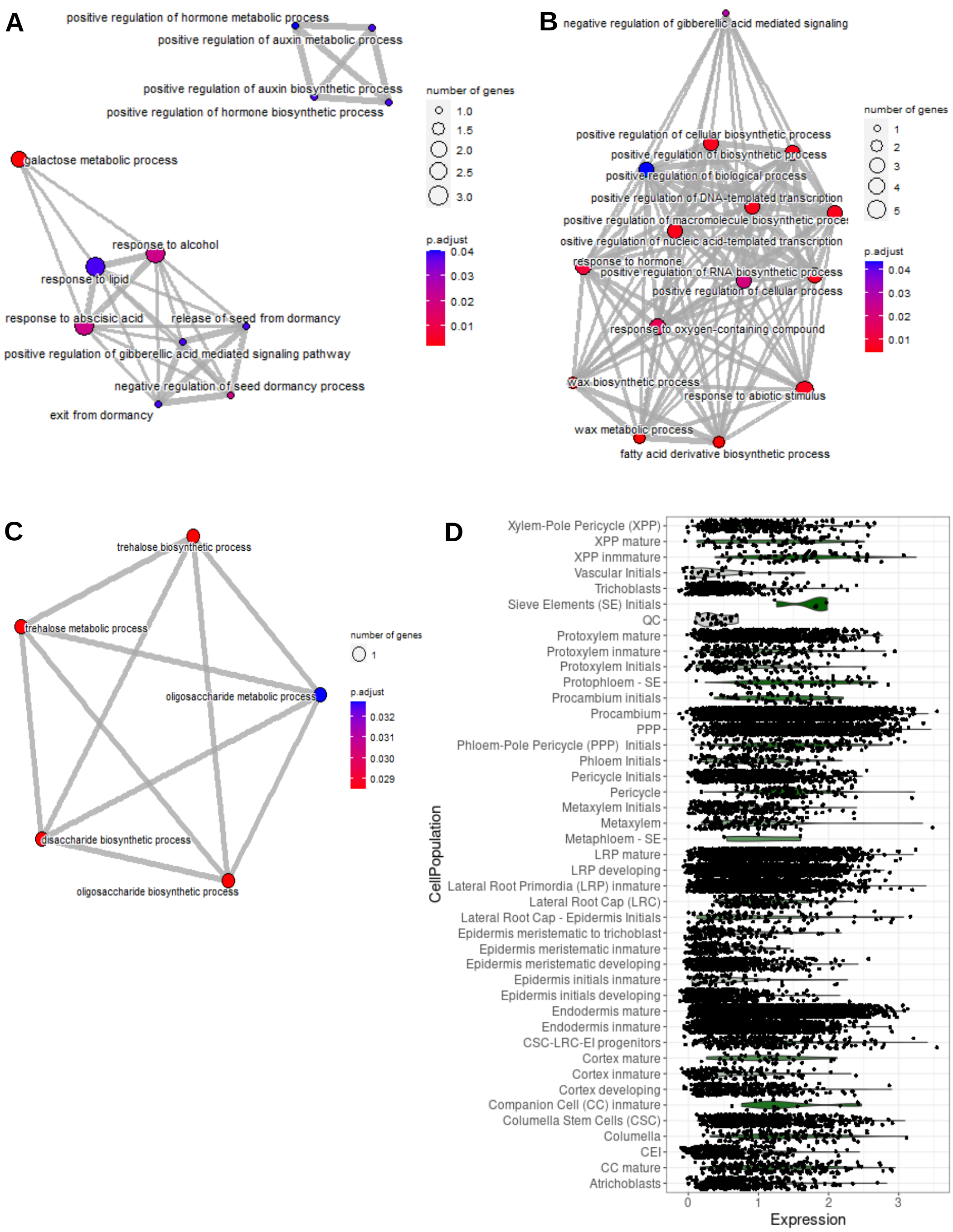


**Fig. S1**. GO term enrichment of genes enriched in **A)** pericycle and **B)** lateral root primordia cell populations of the radial dataset and **C)** phloem pole pericycle (PPP) cell population of single cell experiment. The query genes are upregulated genes (FDR<0.05 and logFC>1.0) in drought conditions, derived from differential expression analysis of BRL3ox vs Col-0 root (Fàbregas et al., 2018). GO terms for biological process with FDR<0.05 are depicted. The size of the dots corresponds to the number of DEGs enriched in the GO term. The color of the dots corresponds to the adjusted p-value (FDR). **D)** Patterns of single cell expression of NAC019, a gene related to drought and enriched in PPP, in all cell populations is shown in the right.

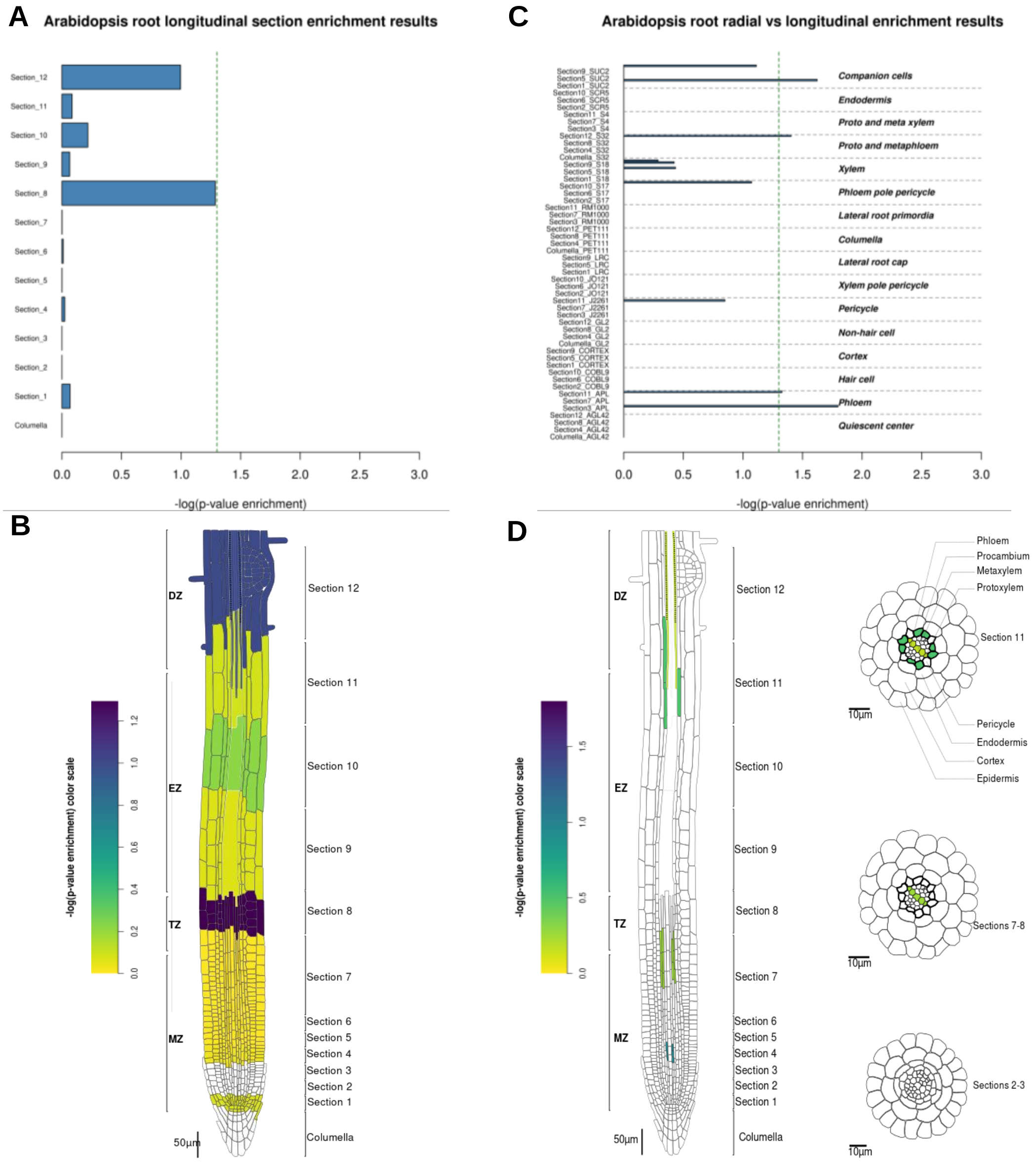


**Fig. S2**. **Validation of TOTEM with bulk transcriptomic data.**

**A)** Longitudinal and **B)** intersected radial-longitudinal expression patterns of upregulated genes in BRL3ox vs Col-0 in Arabidopsis roots (FDR<0.05 and logFC>1.0) under drought conditions. The dashed green vertical line of the barplot correspond to p-value = 0.05.

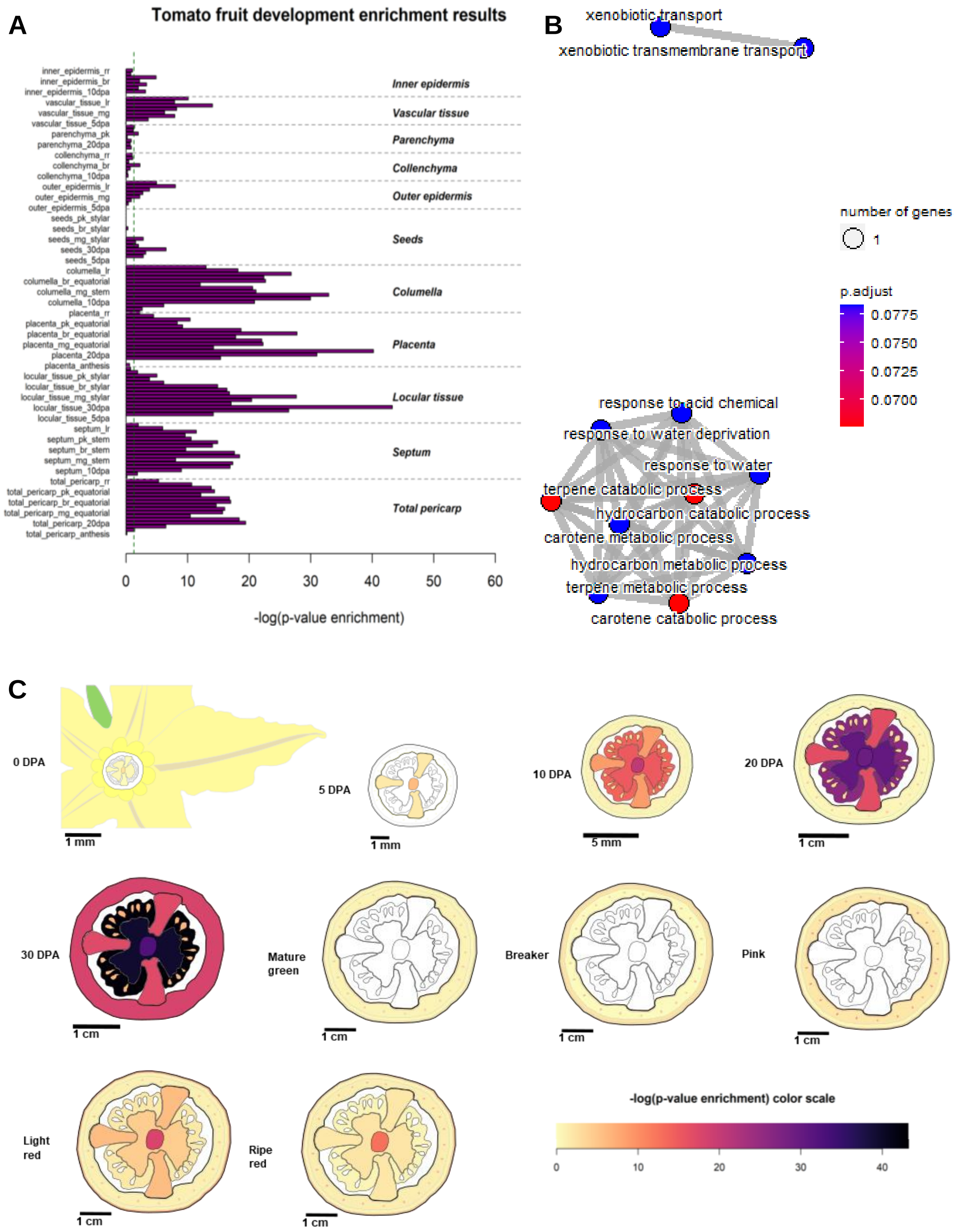


**Fig. S3. Outputs for tomato fruit development based on Shinozaki et al., 2018 experiment.** The query list are drought responsive genes correlated with developmental defects, defined in Lee et al., 2018. GO enrichment of the 139 genes enriched in placenta at 30 DPA is depicted in the top-right. The size of the dots corresponds to the number of DEGs enriched in the GO term. The color of the dots corresponds to the adjusted p-value (FDR). The dashed green vertical line of the barplot correspond to p-value = 0.05.

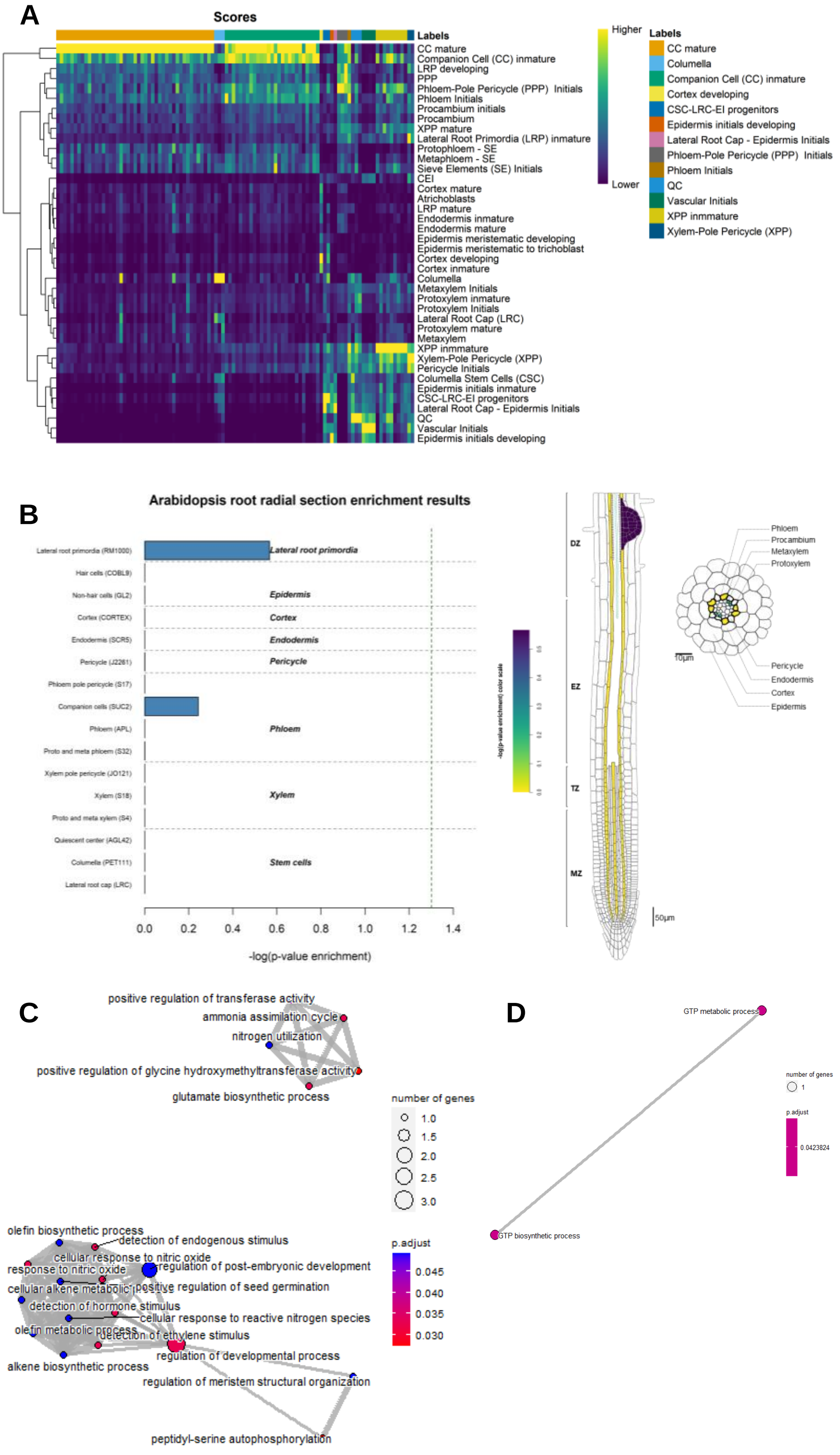


**Fig S4. pSUC2:BRL3:GFP cells annotation onto TOTEM root single cell atlas.**

**A)** SingleR score heatmap for all cells (columns) across all reference labels (rows). Higher scores represented in yellow, lower scores represented in darkblue. **B)** Enrichment pattern in Brady experiment of upregulated genes in pSUC2:BRL3:GFP cells vs WT reference cells (FDR<0.05 and log2(FC) >0.5) represented in form of barplot (left) and vectorial image (right). The dashed green vertical line of the barplot correspond to p-value = 0.05. GO term enrichment of upregulated genes in **C)** Companion cells (SUC2) population and **D**) Lateral root primordia (RM1000).

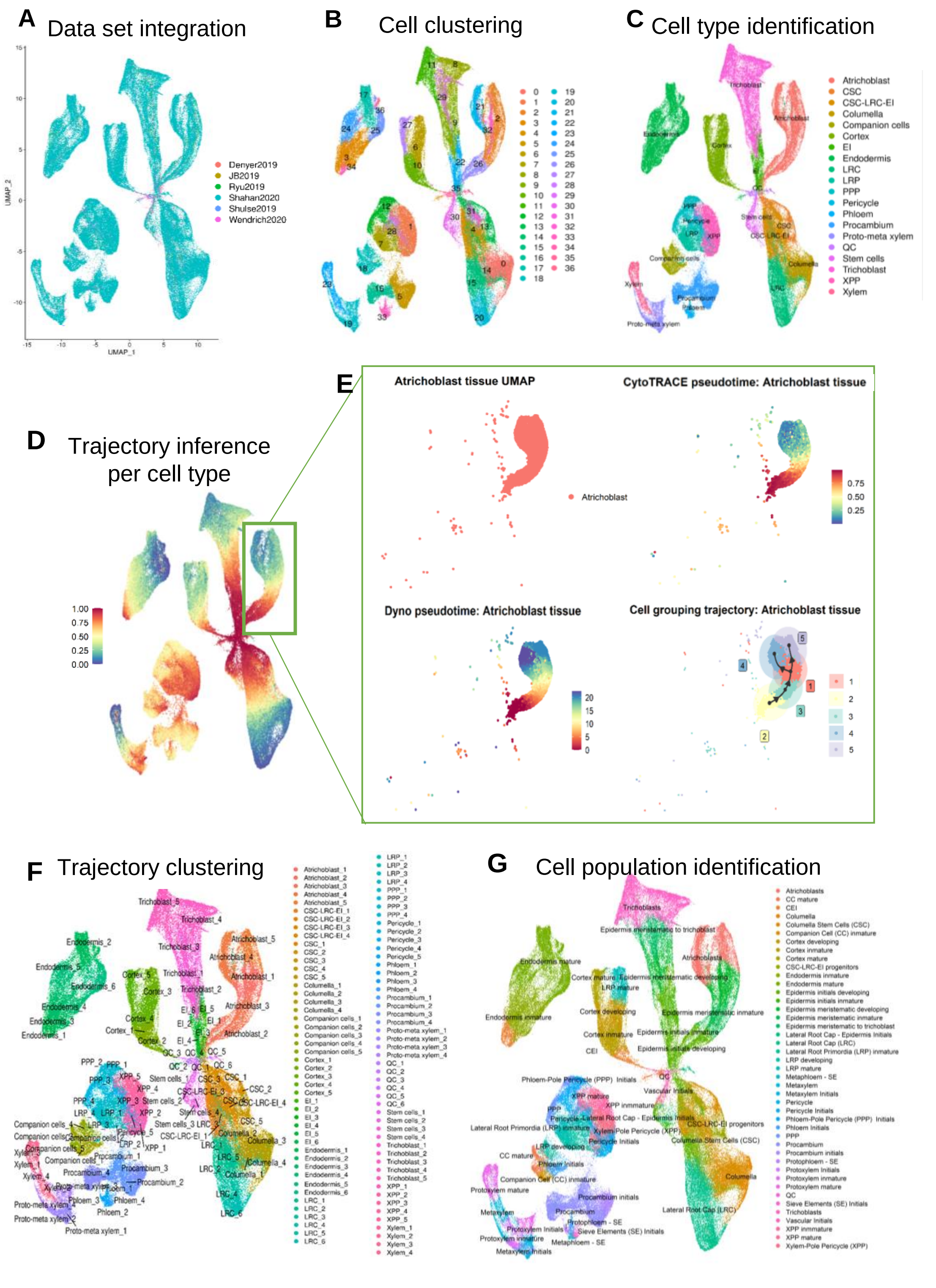


**Fig. S5. Arabidopsis root scRNAseq WT atlas.**

UMAP plot of 154005 cells in root scRNAseq atlas colored and annotated by **A)** the dataset of origin, **B)** shared nearest neighbor (SNN) clustering, **C)** cell type identity and **D)** CytoTRACE pseudotime values. **E)** Atrichoblast trajectory analysis as example: UMAP plot of atrichoblast cells colored and annotated by (top-left) cell type identity, (top-right) CytoTRACE psudotime values, (bottom-left) Dyno pseudotime values, (bottom-right) dyno trajectory groups and trajectory direction. Warmer colours and higher values of CytoTRACE pseudotime values indicate less differenciated cells. Warmer colours and lower values of Dyno pseudotime values indicate less differenciated cells. After trajectory analyses of each cell type, the atlas was annotated by **F)** trajectory groups and **G)** cell population identity.

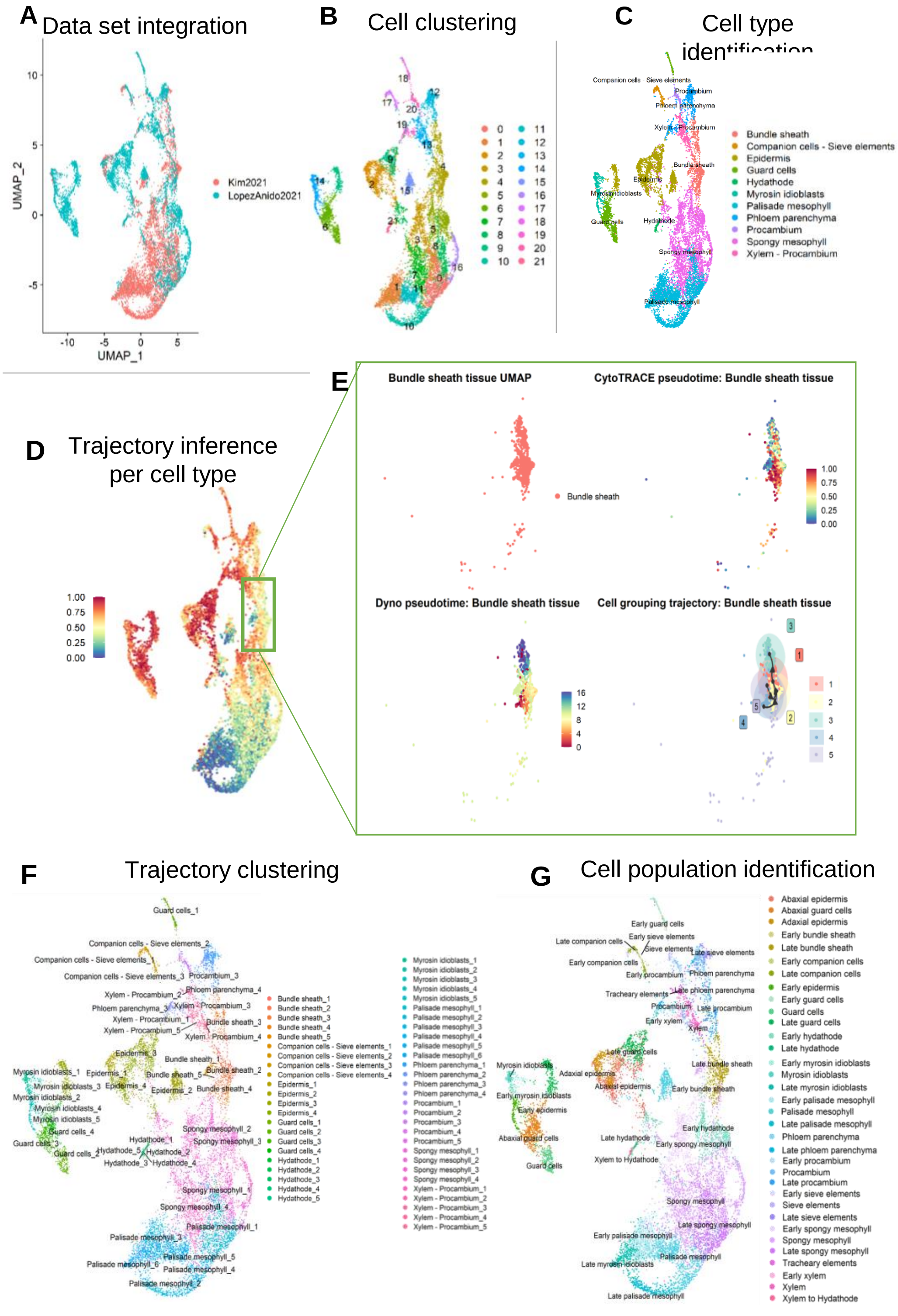


**Fig. S6. Arabidopsis leaf scRNAseq WT atlas.**
UMAP plot of 11413 cells in leaf scRNAseq atlas colored and annotated by **A)** the dataset of origin, **B)** shared nearest neighbor (SNN) clustering, **C)** cell type identity and **D)** CytoTRACE pseudotime values. **E)** Bundle sheath trajectory analysis as example: UMAP plot of bundle sheath cells colored and annotated by (top-left) cell type identity, (top-right) CytoTRACE psudotime values, (bottom-left) Dyno pseudotime values, (bottom-right) dyno trajectory groups and trajectory direction. Warmer colours and higher values of CytoTRACE pseudotime values indicate less differenciated cells. Warmer colours and lower values of Dyno pseudotime values indicate less differenciated cells. After trajectory analyses of each cell type, the atlas was annotated by **F)** trajectory groups and **G)** cell population identity.
